# Transcriptome mining extends the host range of the *Flaviviridae* to non-bilaterians

**DOI:** 10.1101/2022.11.24.517790

**Authors:** Jonathon C.O. Mifsud, Vincenzo A. Costa, Mary E. Petrone, Ezequiel M. Marzinelli, Edward C. Holmes, Erin Harvey

**Affiliations:** Sydney Institute for Infectious Diseases, School of Medical Sciences, The University of Sydney, Sydney, NSW 2006, Australia; School of Life and Environmental Sciences, The University of Sydney, Sydney, NSW 2006, Australia; Sydney Institute of Marine Science, 19 Chowder Bay Rd, Mosman, NSW, 2088 Australia; Singapore Centre for Environmental Life Sciences Engineering, Nanyang Technological University, Singapore, 637551 Singapore

**Keywords:** *Flaviviridae*, *Flavivirus*, *Pestivirus*, *Hepacivirus*, virus discovery, Metazoa, phylogeny

## Abstract

The *Flaviviridae* are a family of positive-sense RNA viruses that include well-documented agents of human disease. Despite their importance and ubiquity, the time-scale of flaviviral evolution is uncertain. An ancient origin, spanning time-scales of millions of years, is supported by their presence in both vertebrates and invertebrates and the identification of a flavivirus-derived endogenous viral element in the peach blossom jellyfish genome (*Craspedacusta Sowerby*, phylum Cnidaria), implying that the flaviviruses arose early in the evolution of the Metazoa. To date, however, no exogenous flavivirus sequences have been identified in these hosts. To help resolve the antiquity of the *Flavivirdae* we mined publicly available transcriptome data across the Metazoa. From this, we expanded the diversity within the family through the identification of 32 novel viral sequences, and extended the host range of the pestiviruses to include amphibians, reptiles, and ray-finned fish. Through cophylogenetic analysis we found cross-species transmission to be the predominate macroevolutionary event across the non-vectored flaviviral genera (median, 68%), including a cross-species transmission event between bats and rodents, although long-term virus-host co-divergence was still a regular occurrence (median, 23%). Notably, we discovered flavivirus-like sequences in basal metazoan species, including the first associated with Cnidaria. This sequence formed a basal lineage to the genus *Flavivirus* and was closer to arthropod and crustacean flaviviruses than those in the tamanavirus group that include a variety of invertebrate and vertebrate viruses. Combined, these data attest an ancient origin of the flaviviruses, close to the emergence of the metazoans 750–800 million years ago.

## 1. Introduction

The *Flaviviridae* are a family of positive-sense single-stranded RNA viruses comprised of the genera *Flavivirus, Pestivirus, Pegivirus*, and *Hepacivirus*. These viruses include well-documented agents of human and livestock disease, including dengue virus, hepatitis C virus, yellow fever virus, Zika virus, and Bovine viral diarrhea virus 1. Reflecting their regular occurrence as pathogens, our understanding of flaviviral biology is necessarily skewed towards a subset of metazoan hosts, particularly those known to experience overt disease or act as reservoirs for these viruses, impeding our ability to understand the evolutionary history of this family. Currently available data suggests that all established genera, with the exception of the genus *Flavivirus*, are vertebrate-infecting viruses and do not require an arthropod vector for transmission (Simmonds et al. 2017).

The genus *Flavivirus* can itself be divided into four groups defined by phylogenetic placement and host range; the (i) mosquito-borne flaviviruses (MBFV), (ii) tick-borne flaviviruses (TBFV), (iii) insect-specific flaviviruses (ISF) and (iv) vertebrate-specific flaviviruses, also known as the “no known vector” (NKV) flaviviruses (Blitvich and Firth 2017; Simmonds et al. 2017). A wide diversity of more divergent “flavi-like” viruses have also been identified, including a group associated with crustaceans and decapods as well as the tamanaviruses (after Tamana bat virus (TABV)), which contains viruses from a broad range of vertebrate and invertebrate species (Costa et al. 2021; Geoghegan et al. 2018; Le Lay et al. 2020; Parry et al. 2019; Price 1978; Shi et al. 2018; Skoge et al. 2018; Soto et al. 2020). Another clade of related flavi-like viruses was recently identified in free-living parasitic flatworms (order Tricladida) (Dheilly et al. 2022).

Metagenomic surveys have identified flaviviral sequences with diverse genome structures, straying from the single 9–13 kb polyprotein that previously appeared to be canonical for the family. This expanded diversity includes a group of novel predominantly arthropod associated viruses – the Jingmenviruses – that are both segmented and perhaps multicomponent (Ladner et al. 2016; Qin et al. 2014; Shi et al. 2016; Simmonds et al. 2017). Metagenomic studies have also expanded the host range of hepaci-, pesti- and pegiviruses in non-mammalian hosts, including the discovery of hepaci-and pegiviruses in birds (Chang et al. 2021; Goldberg et al. 2019; Porter et al. 2020; Zhang et al. 2022), hepaci- and pesti-like viruses in cartilaginous fish (Chondrichthyes) (Shi et al. 2018), and hepaciviruses in reptiles and bony fish (Osteichthyes) (Costa et al. 2022; Porter et al. 2020; Shi et al. 2018).

The identification of flaviviral sequences in marine invertebrate and basal vertebrate lineages has led to suggestions that the evolution of the *Flaviviridae* may follow that of the metazoans through virus-host co-divergence over time-scales of hundreds of millions of years (Bamford et al. 2022; Lensink et al. 2022; Shi et al. 2018). This, in turn, has stimulated questions regarding their host range and mode of transmission, while the complex evolutionary history of the flaviviruses and related sequences has been highlighted by their broad host range and sequence diversity. For example, the large phylogenetic gap between the cartilaginous fish and mammalian pestiviruses suggests that related viruses in bony fish, amphibians, reptiles, and birds exist but have yet to be sampled. The identification of flaviviruses in freshwater and marine crustaceans and a flavivirus-derived endogenous viral element (EVE) in the peach blossom jellyfish genome (*Craspedacusta Sowerby*, phylum Cnidaria) (Bamford et al. 2022) points towards an aquatic origin for the flaviviruses and highlights their long evolutionary association with the Metazoa. In particular, the cnidarian EVE suggests the existence of exogenous cnidarian flaviviruses. These are of importance for understanding the evolution of the *Flaviviridae*, as cnidarians, which include jellyfish, sea anemones, and corals, are an early branching lineage of the metazoans thought to have originated 700 million years ago (Erwin 2015). The phylogeny of the Metazoa can itself be divided into two major groups: those with bilateral body symmetry, the bilaterians, that comprise 99% of all animal species and, basal to them, the non-bilaterians, that include all the early diverging metazoan lineages – the Cnidaria, Placozoa, Porifera, and Ctenophora. Because non-bilaterians lack the body plan and circulatory system of vertebrates, it is possible that viruses of these hosts use an alternate mode of cell-to-cell transmission. To date, however, no flavi-like viruses have been identified in these early diverging metazoan phyla.

Transcriptome mining is a proven method of virus discovery that leverages previous investment in metagenomics (Dheilly et al. 2022; Edgar et al. 2022; Greninger 2018; Grimwood et al. 2021; Iwamoto et al. 2021; Mifsud et al. 2022; Miller et al. 2021; Olendraite et al. 2022; Paraskevopoulou et al. 2021; Parry et al. 2019). To understand the host range of flaviviral sequences throughout the Metazoa and hence more accurately determine the age of the *Flaviviridae*, we used the Serratus RNA-dependent RNA polymerase (RdRp) search (https://www.serratus.io/explorer/rdrp) to mine the Sequence Read Archive (SRA) database for novel flaviviral sequences.

## 2. Methods

### 2.1 Screening of SRAs for flavivirial-like sequences

The Serratus RdRp search and palmID analysis suite (Babaian and Edgar 2022; Edgar et al. 2022) were used to identify data sets within the SRA (as of May, 2022) that contain signatures of novel flavivirus-like sequences. This search was limited to the *Flaviviridae* with a threshold score of >= 50. The *de novo* transcriptome assemblies available at the National Center for Biotechnology Information (NCBI) Transcriptome Shotgun Assembly (TSA) Database (https://www.ncbi.nlm.nih.gov/genbank/tsa/) (as of June, 2021) were also screened using the translated Basic Local Alignment Search Tool (tBLASTn) algorithm under default scoring parameters and the BLOSUM45 matrix. Amino acid sequences from representatives of the four *Flaviviridae* genera along with the related Jingmenviruses were used as queries for the palmID and TSA database searches. The SRA and TSA search range was limited to Eukaryotes (NCBI taxonomic identifier (taxid 2759) excluding the Viridiplantae (taxid 33090). Invertebrate data sets were limited to aquatic species as terrestrial invertebrate SRAs has been previously examined (Paraskevopoulou et al. 2021; Wu et al. 2020).

### 2.2 Tunicate collection, RNA extraction, and metagenomic next-generation sequencing

The tunicate *Botrylloides leachii* was collected by divers wearing surgical gloves at 0.5-3m depth at the pier-pilings in Chowder Bay, Sydney, Australia (site description in Marzinelli (2012)), on 24 November, 2021. Sections of colonies were detached from the substratum using sterile tweezers, which were rinsed in 80% ethanol between samples, and brought to the surface, where they were placed in sterile cryogenic tubes. Samples were stored in liquid nitrogen on site and then transferred to a -80°C freezer until extraction. Total RNA was extracted using the RNeasy Plus Mini Kit (Qiagen, Hilden, Germany) as previously described in (Geoghegan et al. 2021). These libraries were constructed using the Truseq Total RNA Library Preparation Protocol (Illumina). Host ribosomal RNA (rRNA) was depleted with the Ribo-Zero Plus Kit (Illumina) and paired-end sequencing (150 bp) was carried out on the NovaSeq 500 platform (Illlumina). Library construction and metatranscriptomic sequencing were performed by the Australian Genome Research Facility (AGRF).

### 2.2 Identification of novel flavivirial genomes

Raw FASTQ files for all libraries that contained flavivirus-like sequences were obtained through the European Nucleotide Archive (https://www.ebi.ac.uk/ena/browser/home). Adapter removal and quality trimming were conducted using Trimmomatic (v0.38) with parameters SLIDINGWINDOW:4:5, LEADING:5, TRAILING:5 and MINLEN:25 (Bolger et al. 2014). To recover full-length virus sequences, raw reads were assembled *de novo* into contigs using MEGAHIT (v1.2.9) (Li et al. 2015). The assembled contigs were then compared to the NCBI non-redundant protein database (as of August, 2021) and a custom *Flaviviridae* protein database using Diamond BLASTx (v2.0.9) with an e-value threshold of 1×10^−5^ (Buchfink et al. 2015). To identify highly divergent sequences, a custom *Flaviviridae* protein database was regularly updated with the novel viruses identified.

### 2.3 Genome extension and annotation

Sequence reads were mapped onto virus-like contigs using Bbmap (v37.98) and areas of heterogeneous coverage were manually checked using Geneious (v11.0.9) (Bushnell 2014; Kearse et al. 2012). Where possible, the extremities of contigs were manually extended and re-submitted to read mapping until the contig appeared complete or no overhanging extremities were observed. Sequences of vector origin were detected using VecScreen (https://www.ncbi.nlm.nih.gov/tools/vecscreen/) and removed. Contig abundances were calculated using the RNA-Seq by Expectation Maximization (RSEM) software (v1.3.0) (Li and Dewey 2011). GetORF from EMBOSS (v6.6.0) was used to predict open reading frames (ORFs) (Rice et al. 2000). To annotate protein functional domains, the InterProScan software package (v5.56) was used with the TIGRFAMs (v15.0), SFLD (v4.0), PANTHER (v15.0), SuperFamily (v1.75), PROSITE (v2022_01), CDD (v3.18), Pfam (v34.0), SMART (v7.1), PRINTS (v42.0), and CATH-Gene3D databases (v4.3.0) (Jones et al. 2014). Genome diagrams were constructed using a manually curated selection of predicted functional domains and visualized using gggenomes (Thomas Hackl 2022).

### 2.3 Detection of endogenous virus elements

To screen for endogenous viral elements (EVEs) within the viral-like contigs, the putative virus-like nucleotide sequence was compared to the corresponding host genome (where available) and a subset of the whole-genome shotgun (WGS) contig database (as of October 2022) using the TBLASTN algorithm with an E value cutoff of 1 × 10 ^-20^. In addition, the virus-like sequences were checked for host gene contamination using the contamination function implemented in CheckV (v0.8.1) (Nayfach et al. 2021). All EVEs were removed from subsequent analyses.

### 2.4 Assessment of library composition

Taxonomic identification for the contigs assembled for each library was obtained by aligning them to the custom NCBI nt database using the KMA aligner and the CCMetagen program (Clausen et al. 2018; Marcelino et al. 2020). In the case of the cigar comb jelly flavivirus, where raw reads are not publicly available, contigs from the corresponding transcriptome shotgun assembly (TSA: GHXY01000001:GHXY01366104) were used as input. Virus abundance was calculated by counting the number of nucleotides matching the reference sequence with an additional correction for template length (the default parameter in KMA). Krona graphs were created using the KMA and CCMetagen methods and further edited in Adobe Illustrator (https://www.adobe.com) (Clausen et al., 2018; Marcelino et al., 2020).

Virus sequences identified in this study were named using a combination of the host common name—if known—and the associated *Flaviviridae* genera (e.g., harrimaniidae flavivirus). Virus-host assignments were made using a combination of host/virus abundance measurements and phylogenetic analyses. Where <80% of host abundance was associated with the target species of the SRA library, the possibility of alternative hosts was considered. In this case, the other organisms comprising this library were examined to determine if it is possible that they represent the source of the virus sequence. For instance, given the known host range of the flaviviruses it is more likely that these sequences are derived from metazoan species than from bacteria, fungi, or archaea. Therefore, metazoan species were given greater weighting when assigning putative virus-host assignments. Where host assignment proved difficult the suffix “associated” was added to the host name to signify this (e.g., digyalum oweni associated virus).

Where the taxonomic position of a virus was ambiguous, the suffix “-like” was used (e.g., African cichlid flavi-like virus).

### 2.5 Phylogenetic analysis

Phylogenetic trees of the putative flaviviral sequence identified here were inferred using a maximum likelihood approach. Translated virus contigs were aligned with known flaviviral protein sequences from NCBI/GenBank using the program MAFFT (v7.402) with the generalized affine gap cost algorithm (Katoh and Standley 2013; Sayers et al. 2021). Poorly aligned regions were removed using trimAl (v1.2) with a gap threshold ranging from 0.7 to 0.9 and a variable conserve value (Capella-Gutiérrez et al. 2009). All phylogenetic trees were estimated using IQ-TREE2. Branch support was calculated using 1,000 bootstrap replicates with the UFBoot2 algorithm and an implementation of the SH-like approximate likelihood ratio test within IQ-TREE2 (Guindon et al. 2010; Hoang et al. 2017). Selection of the best-fit model of amino acid substitution was determined using the Akaike information criterion (AIC), the corrected AIC, and the Bayesian information criterion with the ModelFinder function in IQ-TREE 2 (Kalyaanamoorthy et al. 2017; Minh et al. 2020). The trimming methods, alignment lengths and phylogenetic models chosen in this analysis are outlined in Supplementary Table 1. Phylogenetic trees were annotated using the R packages phytools (v1.0-3), and ggtree (v3.3.0.9) and further edited in Adobe Illustrator (https://www.adobe.com) (Revell 2012; Yu et al. 2017).

### 2.6 Assessment of cross-species virus transmission

To visualise the relative occurrence of cross-species transmission and virus-host codivergence across the *Flaviviridae* we analysed the cophylogenetic relationship between viruses and their hosts. Host cladograms were created using the phyloT software, a phylogenetic tree generator based on NCBI taxonomy (http://phylot.biobyte.de/). Virus-host associations were obtained from the NCBI virus (Brister et al. 2015; Hatcher et al. 2017) and Virus-Host database (release 213)(Mihara et al. 2016) (accessed 14 September 2022). Tanglegrams that graphically represent the correspondence between host and virus trees were created using the R packages phytools (1.0-3) (Revell 2012) and ape (v5.6-2) (Paradis and Schliep 2019). The virus phylogenies used in the cophylogenies were constructed as detailed above. The relative frequencies of cross-species transmission versus virus-host codivergence were quantified using the Jane package that employs a maximum parsimony approach to establish the best “map” of the virus phylogeny onto the host phylogeny (Conow et al. 2010). The cost of duplication, host-jump, and extinction event types were set to 1.0, while virus-host codivergence was set to zero as it was considered the null event. The number of generations and the population size were set to 100. Jane was chosen over its successor eMPRess (Santichaivekin et al. 2020), as it allows a virus to be associated with multiple host species and handles polytomies (Santichaivekin et al. 2020). For a multi-host virus, each association was represented as a polytomy in the virus phylogeny.

## 3. Results

Screening of transcriptomes within the SRA and TSA databases revealed the presence of flaviviral-like sequences in 154 sequencing libraries. The assembly and mining of these sequencing libraries identified 32 novel virus-like sequences, which were subsequently assigned as hepaci-like (20), flavivirus-like (7), pesti-like (4), and as unclassified flavivirial-like sequences (1) (Table 1, Figure 1). These virus-like sequences were predominately found in metazoan transcriptomes belonging to aquatic species (amphibians, bony fish, cnidarians, comb jellies, crustaceans, and hemichordates) as well as land-dwelling vertebrates (birds, primates, and rodents), while one virus-like sequence was assembled from a non-metazoan, alveolate library. No pegi-like virus sequences were found. We now examine each genus in turn.

**Table 1.**
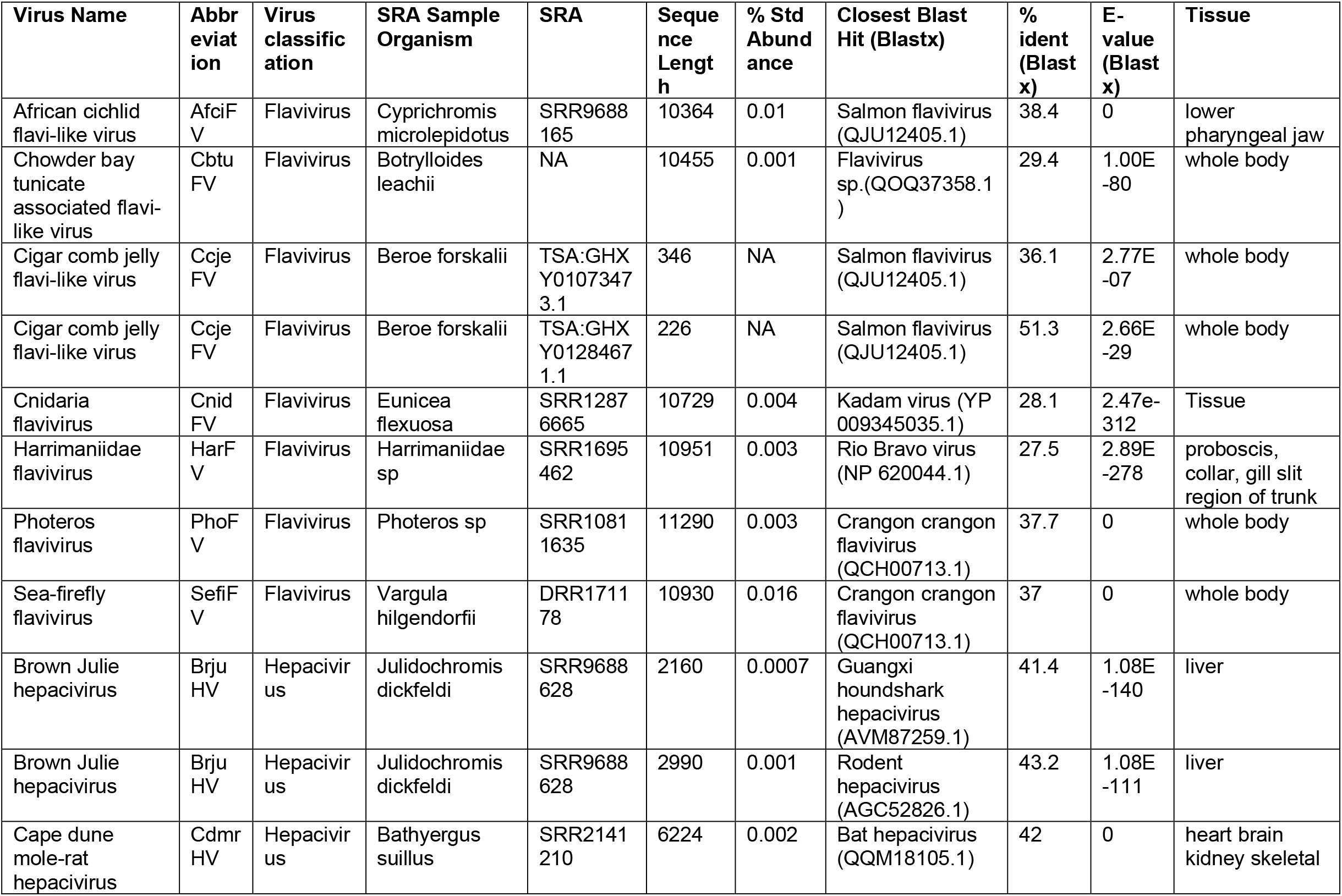
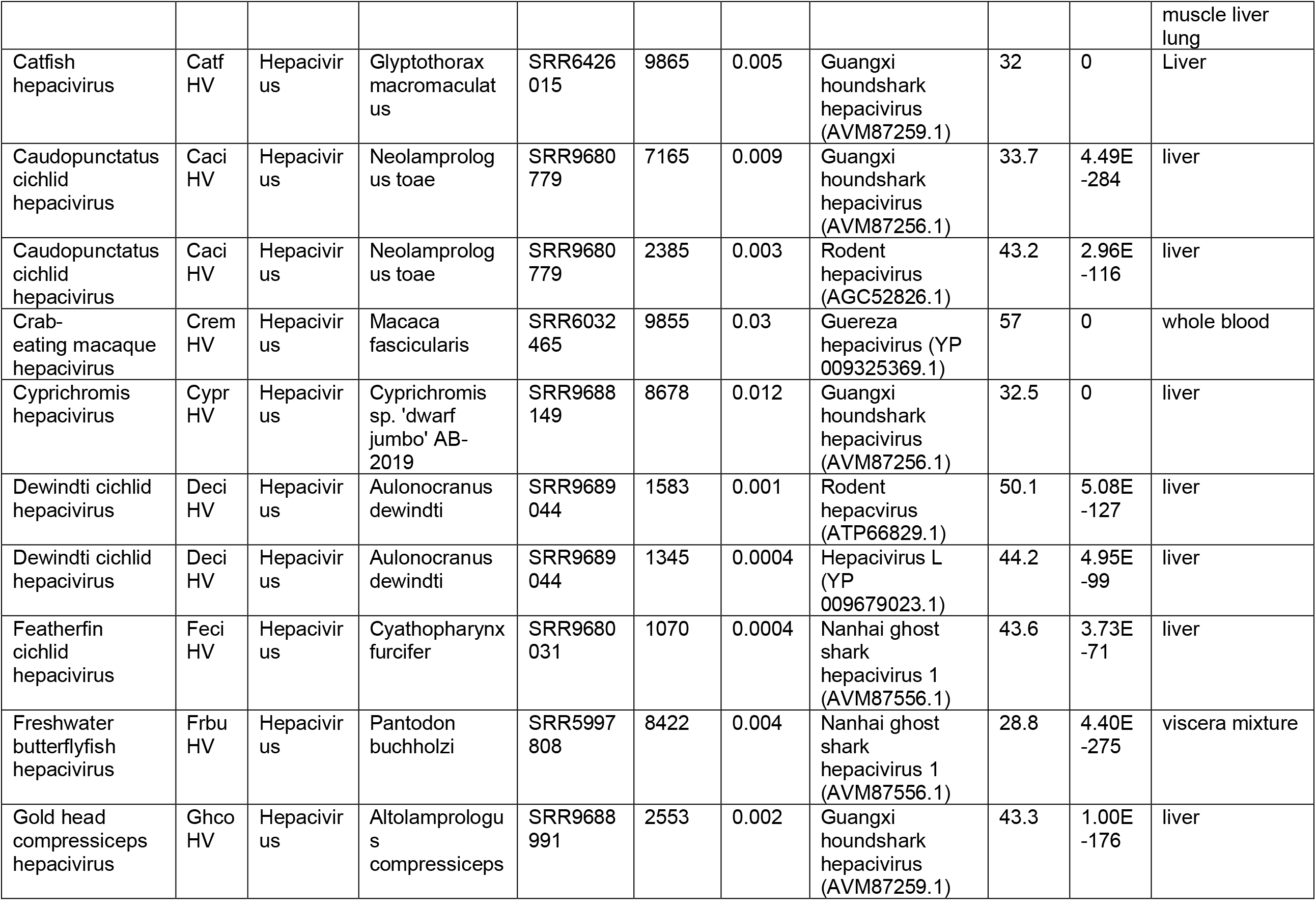
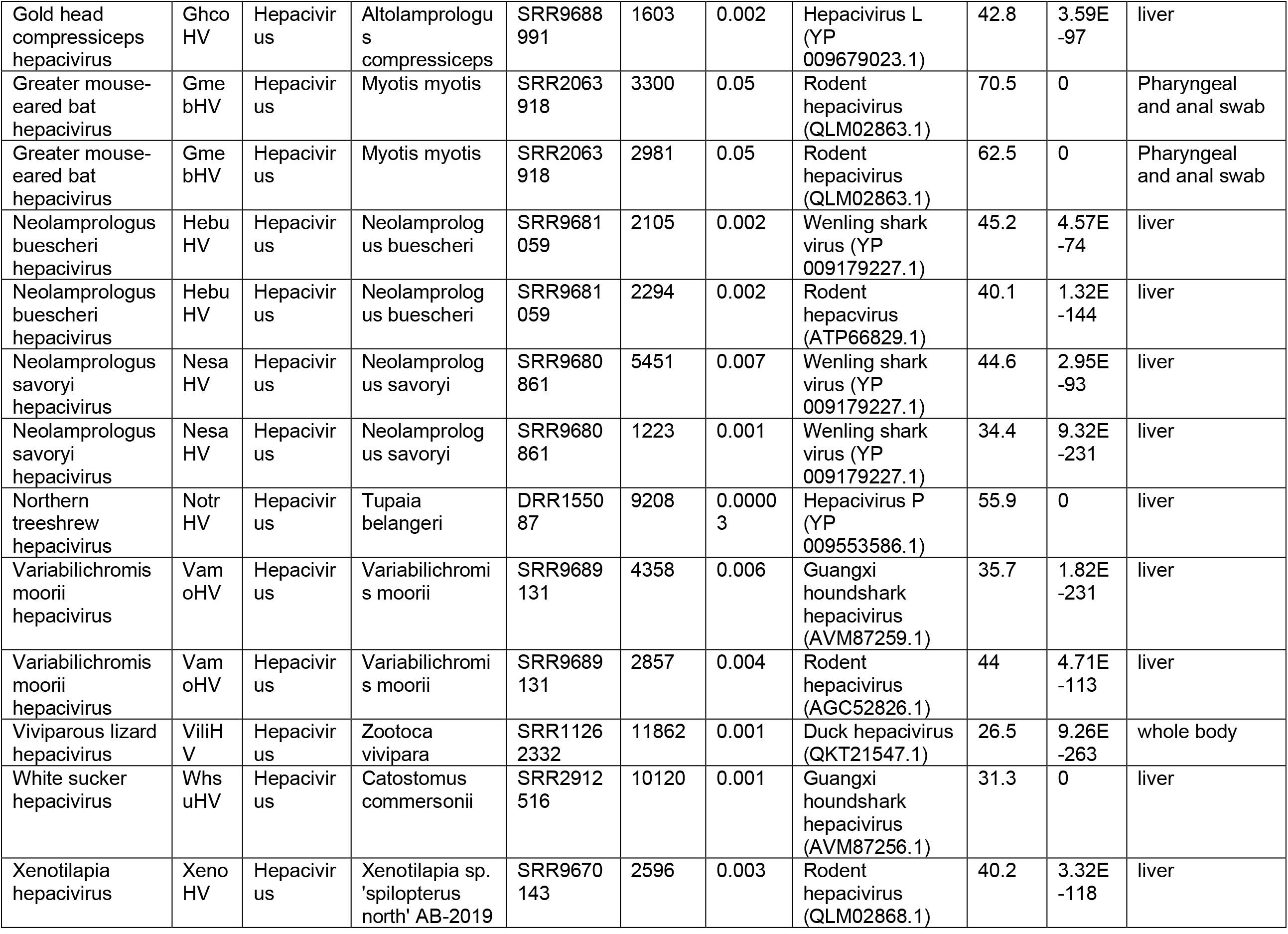
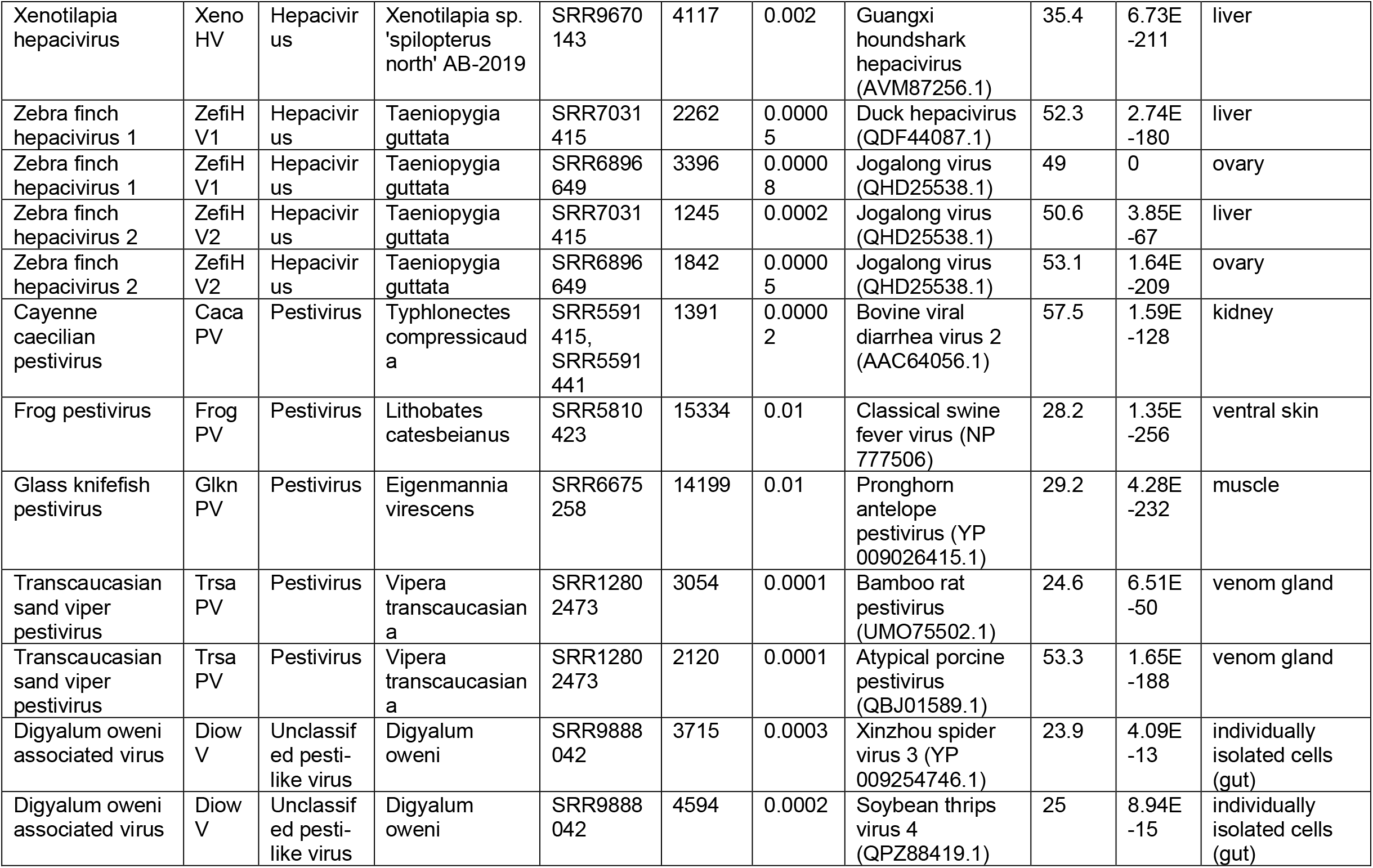
Summary table of the viruses discovered in this study.

**Figure 1.**
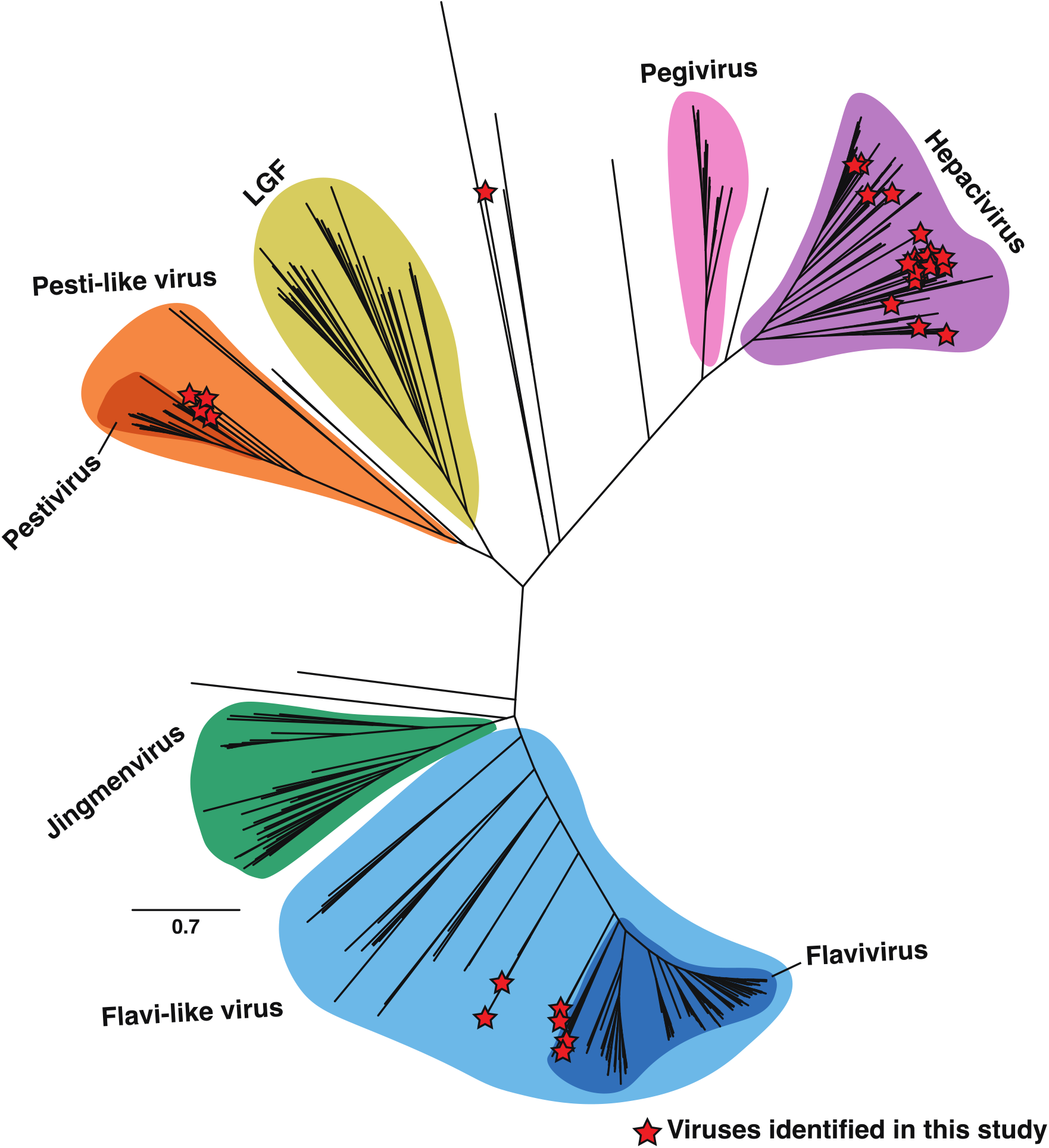
Phylogeny of the *Flaviviridae*. Unrooted maximum likelihood phylogenetic tree of the flaviviral sequences based on the conserved amino acid in the RdRp (NS5). All branches are scaled according to the number of amino acid substitutions per site. Established genera and notable clades that are yet to be ratified by ICTV are highlighted. Novel virus sequences identified in this study are displayed with a red star. LGF refers to the “large genome flaviviruses”.

### 3.1 Genus *Flavivirus*

We identified seven putative flavi-like virus sequences, including cnidaria flavivirus (CnidFV) and cigar comb jelly flavi-like virus (CcjeFV) in libraries of the early diverging metazoan phyla Cnidaria and Ctenophora, harrimaniidae flavivirus (HarFV) in an acorn worm (Enteropneusta), photeros flavivirus (PhoFV) and sea-firefly flavivirus (SefiFV) in marine ostracods, Chowder Bay tunicate-associated flavivirus in tunicates (CbtuFV), and African cichlid flavivirus (AfciFV) in a cichlid fish (Figure 2). For all but one of these sequences (CcjeFV), complete genome sequences ranging in length from 10,364 to 11,290 nucleotides were assembled. CcjeFV consists of two partial RdRp fragments, 346 and 226 bp in length.

**Figure 2.**
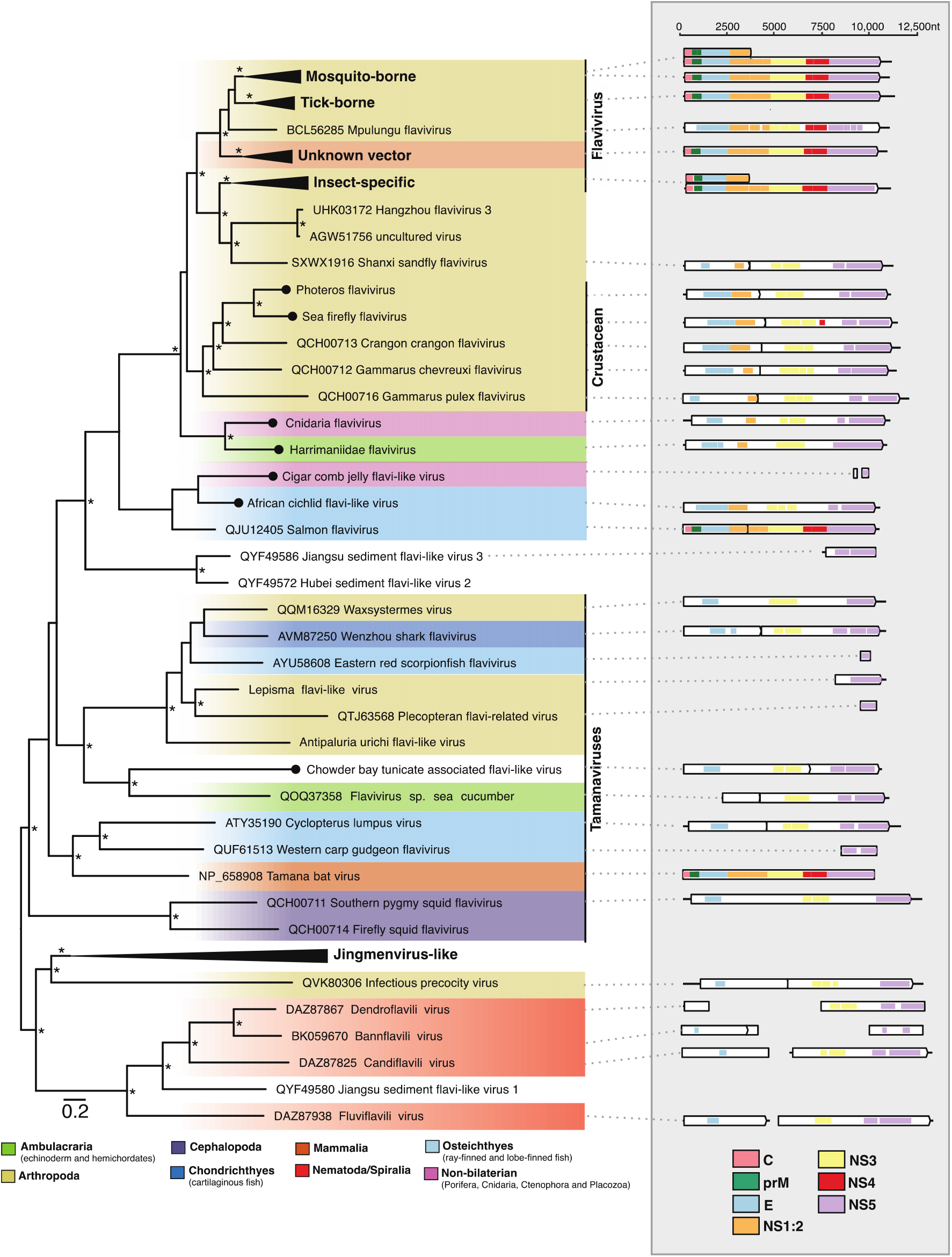
Phylogenetic relationships of the flavi-like viruses identified in this study. (Left) Phylogenetic relationships of the flavi- and jingmenviruses. ML phylogenetic trees based on the conserved amino acid in the RdRp (NS5) show the topological position of virus-like sequences discovered in this study (black circles) in the context of their closest relatives. Branches are highlighted to represent host clade (Ambulacraria = green, Arthropoda = khaki, Cephalopoda = purple, Chondrichthyes = light blue, Mammalia = orange, Nematoda/Spiralia = red, Osteichthyes = dark blue, non-bilaterian = light purple). All branches are scaled to the number of amino acid substitutions per site, and trees were midpoint rooted for clarity only. An asterisk indicates node support where SH-aLRT >= 80% and UFboot >= 95%. (Right) Genomic organization of the virus sequences identified in this study and representative species used in the phylogeny. The data underlying this figure and definitions of acronyms used are presented in Supplementary Table 2.

A range of genome structures were observed and found to be largely consistent with those found in this genus. For example, PhoFV and SefiFV, like the other viruses identified in marine crustaceans, are predicted to contain a programmed −1 ribosomal frameshift on a “slippery” heptanucleotide sequence downstream of the NS1 region (Parry et al. 2019) (Figure 2, Supplementary Figure 1). However, CbtuFV was predicted to contain two ORFs, with the NS4/5 region encoded on the second open reading frame, although no “slippery” heptanucleotide motifs could be detected (Figure 2). The remaining full-length sequences were predicted to contain a single ORF. Virus domains consistent with this genus were detected across all sequences (Figure 2).

Phylogenetic analyses of the conserved NS5 region place the ostracod sequences (PhoFV, SefiFV) within a larger diversity of marine crustacean flaviviruses. Two sequences, CnidFV and HarFV, fell basal to all classified members of the genus *Flavivirus* along with the crustacean flaviviruses (Figure 2). Notably, these sequences appear closer in amino acid identity to tick, insect-specific and crustacean flaviviruses than those in the tamanavirus clade. The flavivirus-derived EVEs identified in the Cnidaria fell in approximately the same phylogenetic location as CnidFV and SefiFV (Supplementary Figure 2). CcjeFV and AfciFV were placed phylogenetically with salmon flavivirus (QJU12405.1), although unlike salmon flavivirus AfciFV consists of a single ORF.

### 3.2 Genus *Pestivirus*

We identified four pesti-like virus sequences in amphibians, reptiles, and bony fish (Table 1). Two full genomes, glass knifefish pestivirus (GlknPV) and frog pestivirus (FrogPV) were recovered, ranging from 14199 to 15334bp in length, in addition to two partial genomes, Transcaucasian sand viper pestivirus (FrogPV), and Cayenne caecilian pestivirus (CacaPV) (Figure 3). These sequences exhibit more sequence similarity with mammalian pestiviruses than those associated with cartilaginous fish, with an average of 28% versus 24% amino acid identity across the complete polyprotein. This is reflected in the phylogenetic positioning of the novel pesti-like viruses based on the conserved NS5 region (Figure 3). The newly identified reptile and amphibian pesti-like virus sequences, FrogPV and CacaPV, form a sister group to those found in rodents, bats, and pigs, while the sequence discovered in fish, GlknPV, fell basal to this group but remained as a sister group to those viruses from cartilaginous fish (Shi et al. 2018). The topology of the pestivirus phylogeny varied depending on whether the NS3 or NS5 domains were used to create the alignment. In particular, FrogPV formed a sister lineage to the known pestiviruses in a phylogeny based upon the NS3 region (Figure 3, Supplementary Figure 3).

**Figure 3.**
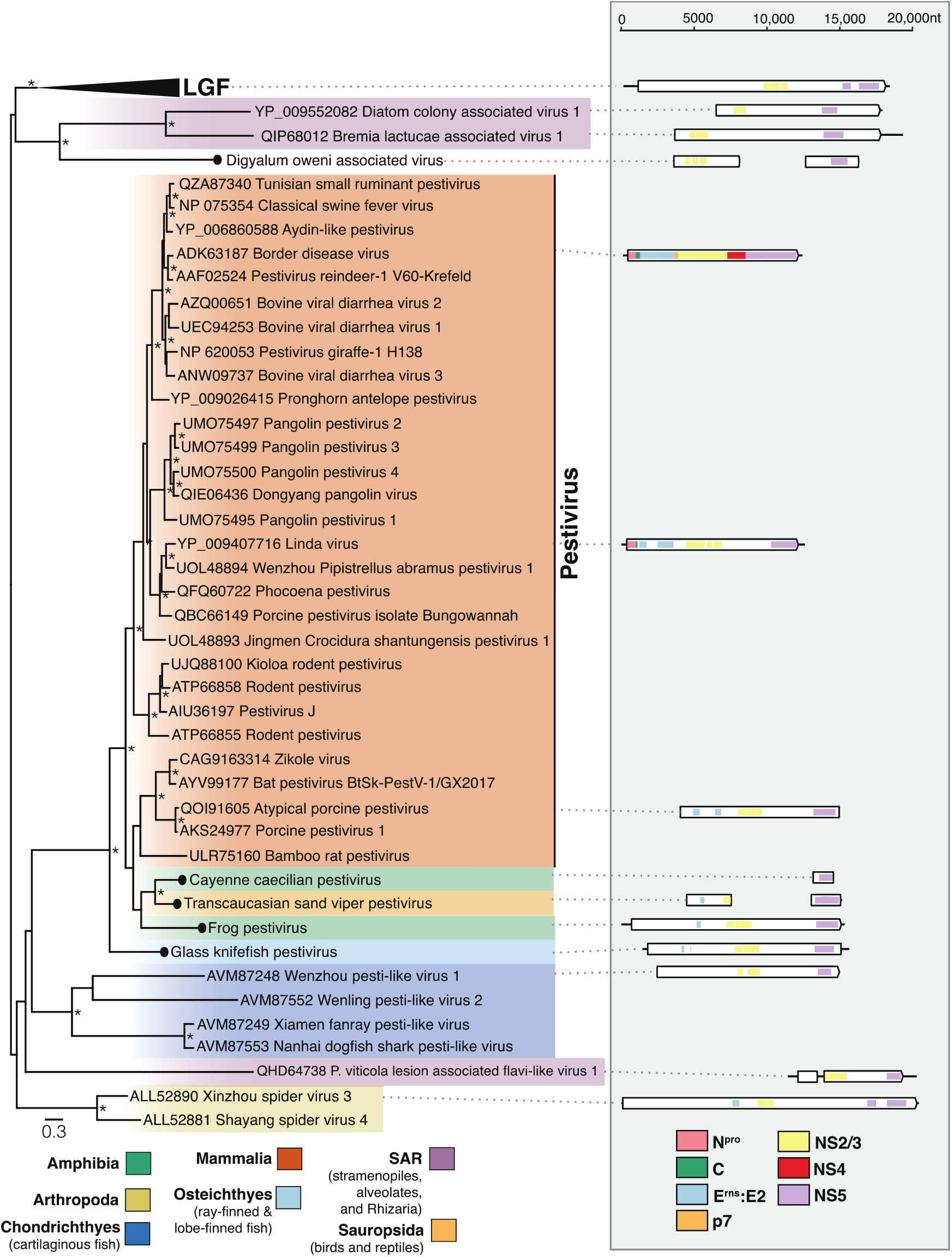
Phylogenetic relationships of the pesti-like viruses identified in this study. (Left) Phylogenetic relationships of the pestiviruses and unclassified relatives. ML phylogenetic trees based on the conserved amino acid in the RdRp (NS5) show the topological position of virus-like sequences discovered in this study (black circles) in the context of their closest relatives. The colour scheme is as found in Figure 2, with the following exceptions, Amphibia = green, Sauropsida = light orange, SAR = light purple. All branches are scaled to the number of amino acid substitutions per site, and trees were midpoint rooted for clarity only. An asterisk indicates node support where SH-aLRT >= 80% and UFboot >= 95%. LGF refers to the “large genome flaviviruses”. Non-novel sequences without NCBI accession were obtained from Wu et al. (2020). (Right) Genomic organization of the virus sequences identified in this study and representative species used in the phylogeny. The data underlying this figure and definitions of acronyms used are presented in Supplementary Table 2.

### 3.3 Genus *Hepacivirus*

We identified 20 novel hepacivirus sequences, of which 14 were found in ray-finned fish (Actinopterygii), expanding on the two hepaciviruses previously identified in this group (Figure 4). The remaining sequences (n=6) add to the known diversity of bat, avian, primate, rodent, and treeshrew hepaciviruses (Figure 4). Of the novel hepaciviruses, five complete genomes were assembled, ranging from 9,208 to 11,862bp in length (Figure 4). Partial genomes sequences containing at least the NS3 and NS5 domains were assembled for the remaining sequences apart from featherfin cichlid hepacivirus, for which only the NS5 region could be assembled (Figure 4). Of note, greater mouse-eared bat hepacivirus (GmebHV) was assembled from a library generated for the analysis of bat viromes (Wu et al. 2012) and shares 70% amino acid identity with rodent hepacivirus (QLM02863.1).

**Figure 4.**
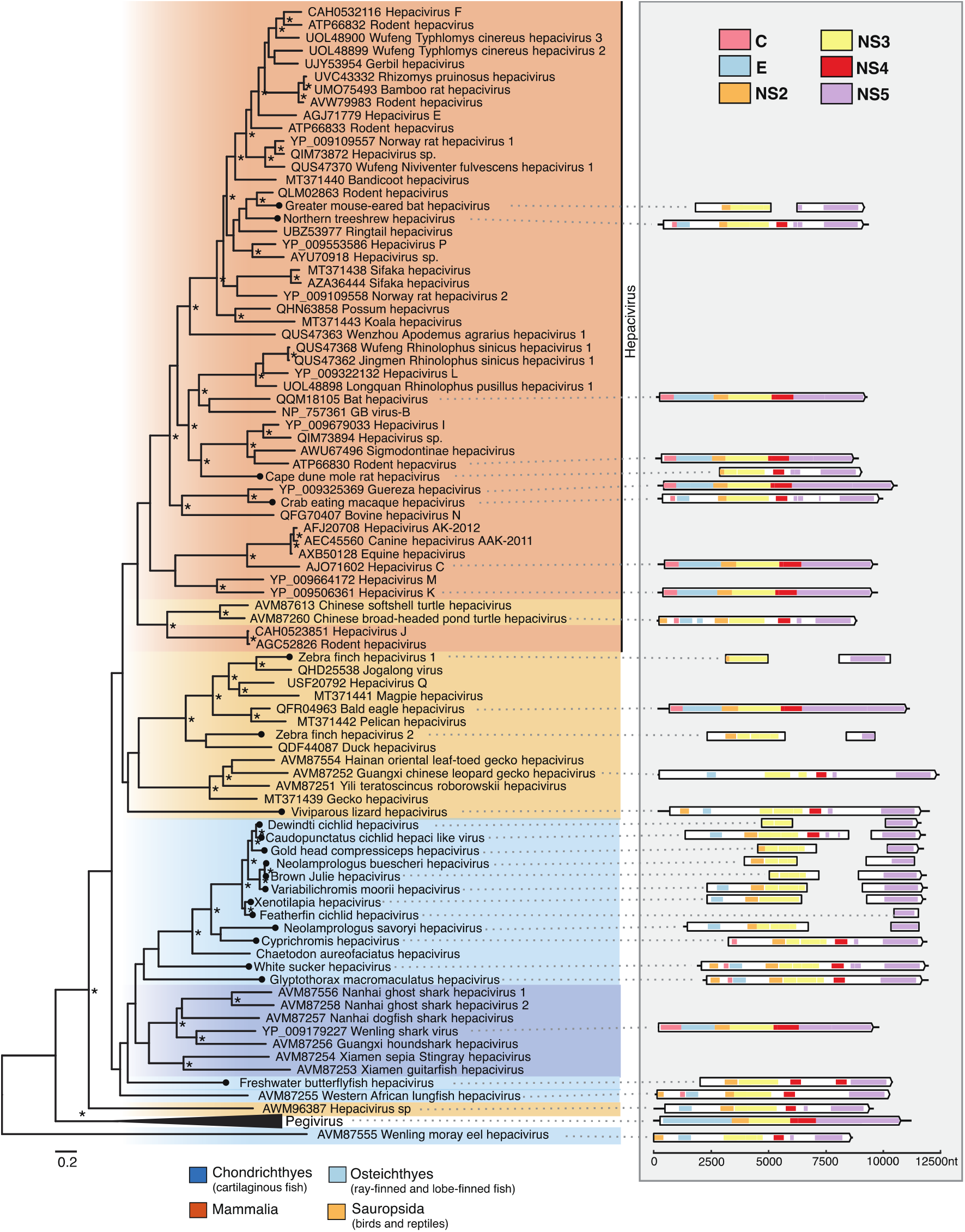
Phylogenetic relationships of the hepaciviruses viruses identified in this study. (Left) Phylogenetic relationships of the ‘pegi-hepaci’ clade. ML phylogenetic trees based on the conserved amino acid in the RdRp (NS5) show the topological position of virus-like sequences discovered in this study (black circles) in the context of their closest relatives. The colour scheme is as found in Figure 2, with the following exception, Sauropsida = light orange. All branches are scaled to the number of amino acid substitutions per site, and trees were midpoint rooted for clarity only. An asterisk indicates node support where SH-aLRT >= 80% and Ufboot >= 95%. (Right) Genomic organization of the virus sequences identified in this study and representative species used in the phylogeny. The data underlying this figure and definitions of acronyms used are presented in Supplementary Table 2.

### 3.4 An unclassified flaviviral-like virus

In addition to the viruses that fell within established genera, we identified a partial flavi-like virus sequence termed digyalum oweni associated virus (DiowV) within the library of *Digyalum oweni*, a species of parasitic protist belonging to the phylum Apicomplexa. Two contigs were assembled from this library, 3689 and 4577bp in length and predicted to contain the NS3 and NS5 domains, respectively (Figure 3). DiowV shares the greatest sequence similarity with the Xinzhou spider virus 3 (YP_009254746) among other large genome flaviviruses (LGF). When included in the ‘pesti-LGF’ tree, DiowV, along with diatom colony associated virus 1 (YP_009552082) and bremia lactucae associated virus 1 (QIP68012) forms a sister group to the LGF. However, in the family-wide tree, these sequences, along with Snake River alfalfa virus (ON669064) fall outside of the ‘pesti-LGF’ lineage, basal to the ‘pegi-hepaci’ group, although these branches receive poor bootstrap support (Figure 1).

### 3.5 Genetic composition of sequencing libraries

Metagenomic sequencing libraries are often comprised of organisms in addition to the target host, which can complicate virus-host assignment. To quantify the composition of these libraries and improve virus host assignments we utilised the KMA and CCMetagen tools (Figure 5). For 20 of the libraries, over 80% of eukaryotic contigs were assigned to the target of the sequencing library (median, 90%; range, 0% to 98%). *Symbiodinium* species represented 64% of all contigs in the *E. flexuosa* (family *Plexauridae*) library in which CnidFV was assembled (Figure 5). In this library, soft corals (order Alcyonacea) represented 63% of metazoan abundance, while tunicates and bony fish represented 13% and 10% of abundance, respectively. Despite *Plexauridae* comprising 60% of cnidarian abundance, other cnidarian families were also detected, including *Ellisellidae, Nephtheidae, Acanthogorgiidae*, and *Nidaliidae*, each representing ∼10% of cnidarian abundance. Likewise, the tunicate library from which CbtuFV was assembled is composed of reads belonging to various marine organisms, including Bryozoa, Cnidaria, and crustaceans, representing an average of 8% abundance each.

**Figure 5.**
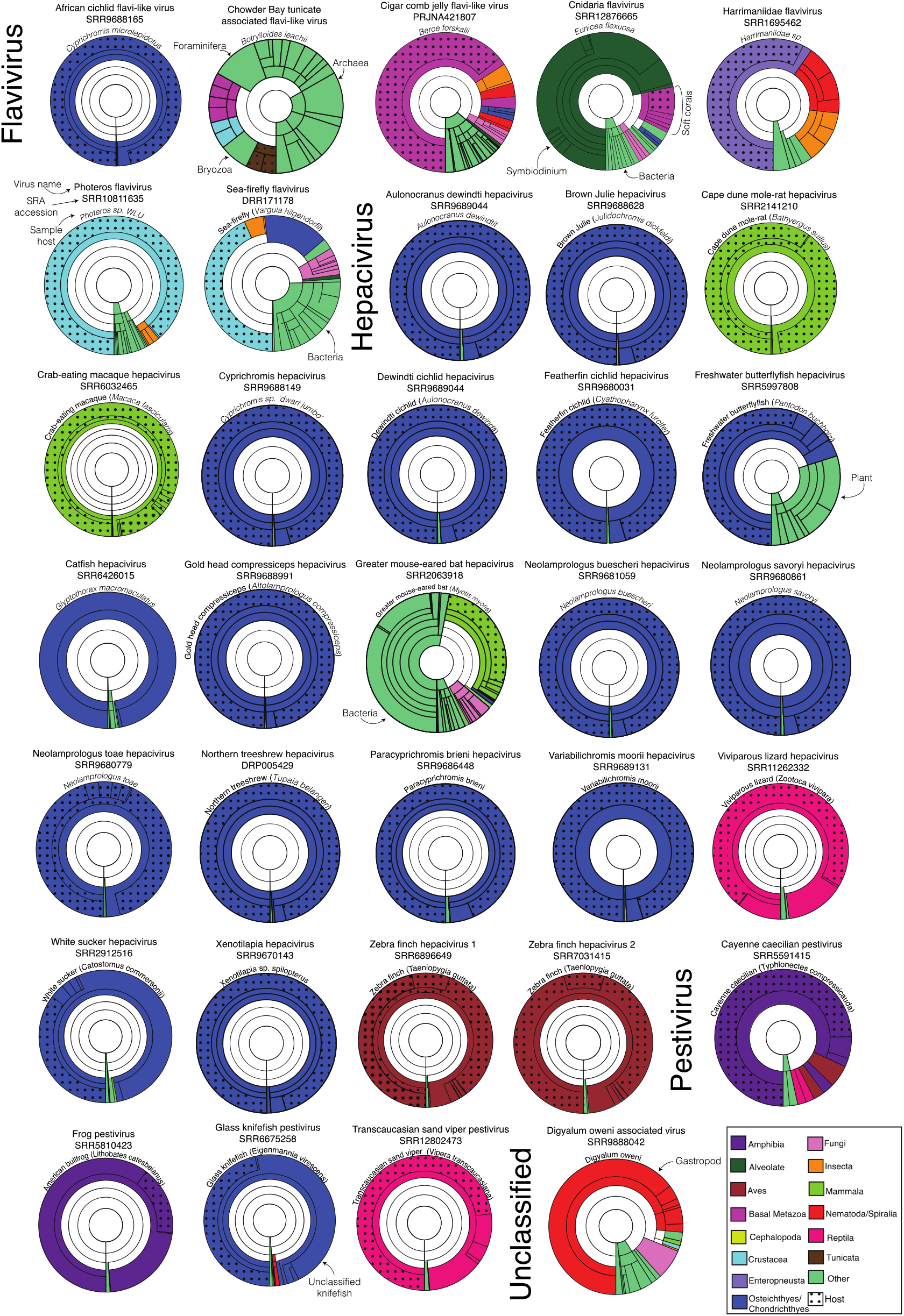
Taxonomic assignments of contigs in sequencing libraries. Each Krona graph illustrates the relative abundance of taxa in a metatranscriptome at varying taxonomic levels. For clarity, a maximum depth of five taxonomic levels was chosen for each graph. The library Sequence Read Archive accession number, host species, and the corresponding virus of interest are annotated above each graph. Segments are highlighted based on the species’ taxonomic grouping. Dots have been used to signify where contigs have been taxonomically assigned within the same family as the host species. Contigs without any matches in the database are not shown.

Contigs belonging to catfish (order Siluriformes) comprised 95% of the *Glyptothorax macromaculatus* library from which catfish hepacivirus (CatfHV) was assembled, but it is not certain to which family of catfish this sample belonged. Likewise, the American bullfrog (*Lithobates catesbeianus)* transcriptome comprised 60% contigs associated with fork-tongued frogs (*Dicroglossidae*) and 17% associated with true frogs (*Ranidae*), including *L. catesbeianus*. No host-associated contigs were detected in the *Digyalum oweni* library in which DiowV was assembled. Instead, 64% of the library is composed of contigs associated with marine gastropod molluscs.

### 3.6 Long-term virus-host evolutionary relationships

To examine the frequency of four macroevolutionary events (i.e., co-divergence, duplication, host-switching, and extinction) among the *Flaviviridae*, we estimated cophylogenies displaying the evolutionary relationship between the flaviviral genera and their hosts (Figure 6). This revealed that cross-species transmission was the most common evolutionary event across the ‘pegi-hepaci’ and pestivirus clades, representing 65% and 71% of events, respectively (Supplementary Figure 4). Two viruses, GmebHV and freshwater butterflyfish hepacivirus (FrbuHV), identified in this study present notable exceptions to this (Figure 6). GmebHV is distinct from known bat hepaciviruses (Hepacivirus K, Hepacivirus L and Hepacivirus M), and instead, groups with those found in rodents, shrews, sloths, and raccoons (Figure 4). FrbuHV, along with Western African lungfish hepacivirus and Wenling moray eel hepacivirus, fall basal to those identified in cartilaginous fish.

**Figure 6.**
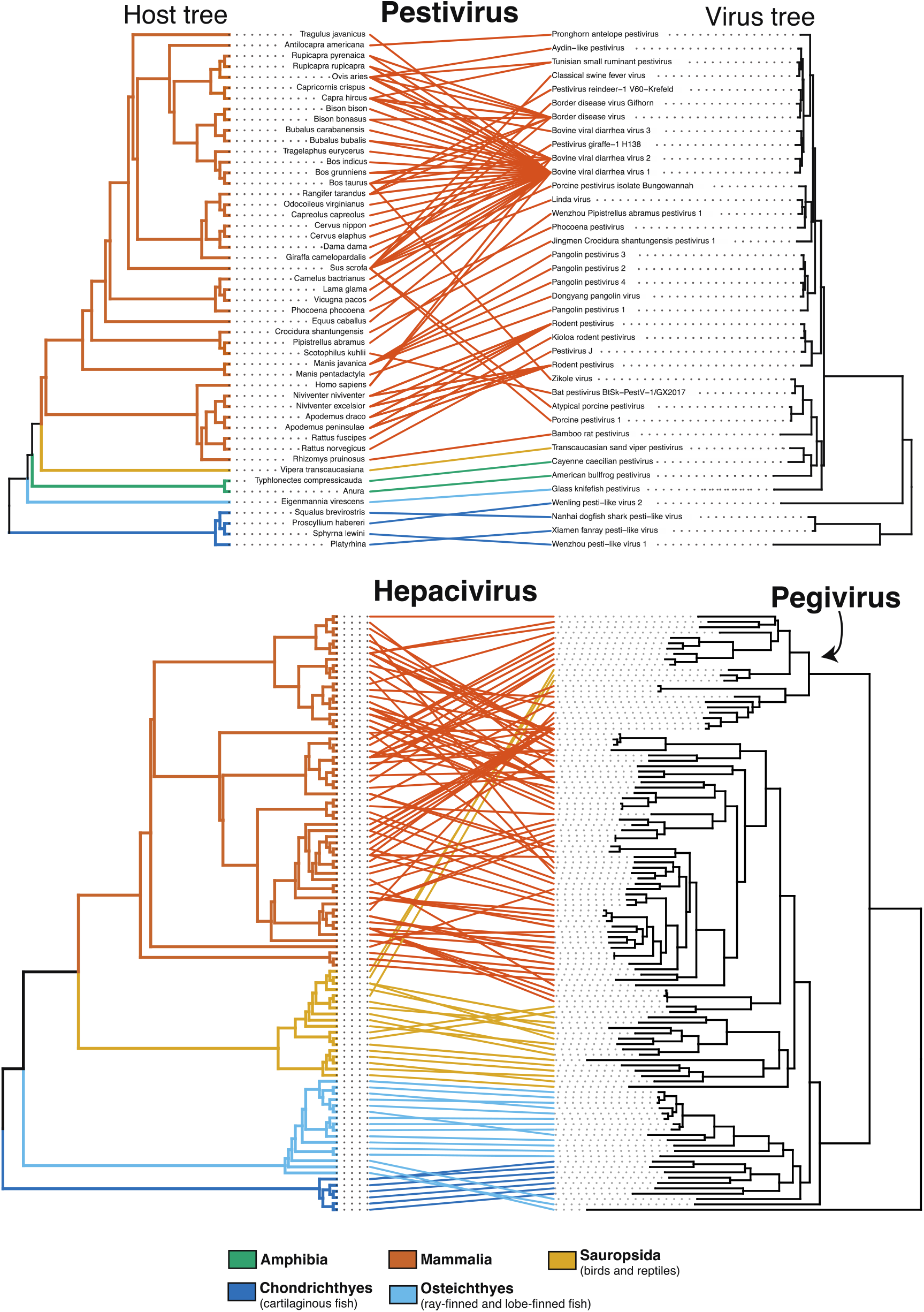
Tanglegram of rooted phylogenetic trees for representative virus groups and their hosts. Branches of the host tree (left) and lines are colored to represent the host clade. The colour scheme is as found in Figure 2, with the following exceptions, Amphibia = green, Sauropsida = light orange. All branches on the virus tree are scaled to the number of amino acid substitutions per site, and both trees were midpoint rooted for clarity only. Supplementary Figure 5 provides the names of the hosts and viruses for the ‘pegi-hepaci’ cophylogeny.

Importantly, despite the widespread occurrence of cross-species transmission, which is expected among the *Flaviviridae* (Geoghegan et al. 2017), virus-host co-divergence was also predicted to have occurred relatively frequently across the ‘pegi-hepaci’ and Pestivirus clades, representing 22% and 23% of all events, respectively. For these groups, duplication events were more uncommon, representing 10% and 6% of total events. Extinction events were rarely predicted, representing 4% of events in the ‘pegi-hepaci’ clade, while no extinction events were detected in the Pestivirus cophylogeny.

## 4. Discussion

Through transcriptome mining we identified 32 novel flaviviral sequences across the Metazoa, including the first flavivirus-like sequences in non-bilaterians, pestivirus-like sequences in amphibians, reptiles, and bony fish, as well as a range of vertebrate hepaciviruses. Hence, this work provides further evidence of the long-term associations between the *Flaviviridae* and Metazoa and highlights the vast number of viruses that remain undiscovered.

The Cnidaria are a primitive and basal metazoan phylum. Through the identification of a flavivirus-like sequence in a cnidarian sample (CnidFV) we suggest that the origins of this group of viruses likely extends much further back in time than previous estimates and closer to the emergence of the metazoans 750–800 million years ago (Erwin 2015). This conclusion is supported by the earlier finding of a flavivirus-derived EVE in the Cnidaria (Bamford et al. 2022). Notably, CnidFV and the cnidrian EVE are more closely related to members of the genus *Flavivirus* than the tamana/flavi-like viruses, suggesting that these groups, including the jingmenviruses, are evolutionary distinct (Bamford et al. 2022). As such, we suggest that the tamana/flavi-like viruses should be given a distinct taxonomic classification within the *Flaviviridae*. Overall, it is difficult to fully resolve the evolutionary history of the flaviviruses with our current understanding of their diversity, although it appears that the origins of this group lie in aquatic environments.

It is important to note that host assignment of the non-bilaterian flaviviruses is tentative as these sequences are extremely divergent and they have been only rarely sampled. Due to the detection of several cnidarian species in addition to the target species, the octocoral *E. flexuosa* in library SRR12876665, we have assigned the resulting virus sequence as cnidaria flavivirus (CnidFV). The high abundance of Symbiodinium in this library is unsurprising given that octocoral-Symbiodinium mutualism is well known (van de Water et al. 2018). However, the phylogenetic placement of this virus with those found in a marine acorn worm suggests it is more likely associated with *E. flexuosa* than Symbiodinium. While CnidFV and the EVE are relatively closely related to each other, there is a large amount of divergence between these sequences. This has been previously observed with crocidura pestivirus and a Crocidura EVE and may reflect divergent evolution since the historic endogenization event (Li et al. 2022).

The discovery of Wenzhou pesti-like virus 1, Wenling pesti-like virus 2, Xiamen fanray pesti-like virus and Nanhai dogfish shark pesti-like virus in cartilaginous fish marked the expansion of the pestiviruses from warm-blooded mammals to basal vertebrate species, suggesting that these viruses infect a range of vertebrate lineages (Shi et al. 2018). For the first time, we identified pesti-like viruses in reptiles, amphibians, and bony fish, extending the host range of these viruses to encompass all vertebrate classes with the exception of Aves. The deep evolutionary association between pestiviruses and vertebrates is further reflected in the clear pattern of pestivirus-host codivergence among the viruses identified in this study. As a result, we anticipate that novel pestiviruses will be found infecting a wider diversity of vertebrates and that their known host range largely reflects where sampling efforts have been directed to date.

Additionally, frog pestivirus was identified in the ventral skin of the American bullfrog, although other species of frog were detected in this library. Within the study in which this library was generated, the American bullfrog appeared resistant to the fungal pathogen *Batrachochytrium dendrobatidis* (Bd) (Eskew et al. 2018). Co-infection with Bd and ranaviruses is frequently observed in frogs, but whether the interactions between these pathogens are antagonistic or facilitative is currently unclear (Bosch et al. 2020). If Bd and pestiviruses are found to commonly co-infect frogs, future efforts should be directed towards studying their interactions.

We identified 20 novel hepacivirus sequences, among which a clade of cichlid-associated hepacivirus sequences is notable. This clade was derived from a single study conducted in the African Great Lake Tanganyika, a freshwater lake shared by Tanzania, the Democratic Republic of the Congo, Burundi, and Zambia that is known for its high diversity of endemic cichlid species (El Taher et al. 2021; Koblmüller et al. 2008). Importantly, the fish and reptile hepaciviruses identified in this study were predominately associated with samples of liver tissue, suggesting hepatotropism is likely a universal feature of these viruses across vertebrates (Smith et al. 2016).

Bats and rodents harbour a large diversity of hepaciviruses and are thought to have played an important role in their global spread and broader evolutionary history (Bletsa et al. 2021; de Souza et al. 2019; Drexler et al. 2013; Epstein et al. 2010; Kapoor et al. 2013; Quan et al. 2013). We identified GmebHV, which falls within a clade of rodent, sloth, and raccoon hepaciviruses. The clear relatedness between GmebHV and rodent hepacivirus (QLM02863), combined with evidence from our co-evolutionary analyses, suggests that this sequence might represent a cross-species transmission event between bats and rodents. Similarly, ancestral state reconstructions have previously shown that cross-species transmission from rodents is likely the source of the sloth and ringtail hepaciviruses (Jo et al. 2022; Moreira-Soto et al. 2020). In this case, we cannot resolve the direction of virus transmission with any certainty or whether intermediate species are involved.

In broad terms, we find that the hepaci-, pesti-, and pegiviruses cluster with the phylogeny of their hosts. Within these groups, cross-species transmission events were limited to within host class (i.e., Mammalia, Sauropsida and Chondrichthyes) except for FrbuHV, Western African lungfish hepacivirus, and Wenling moray eel hepacivirus, which fall basal to those identified in cartilaginous fish. However, the inferred position of these viruses should be treated with caution as these nodes have weak bootstrap support (Figure 4, Figure 6). The clear phylogenetically defined barriers between host classes may reflect differences in receptor binding and cell entry mechanisms among distantly related hosts (Parrish et al. 2008). Host ecology also likely contributes to these barriers, particularly as fewer cross species transmission events are expected to occur between marine and land vertebrates than among land vertebrates, even if receptor binding and cell entry methods were compatible (French et al. 2022; Luis et al. 2015). At deeper taxonomic levels, we observed clear patterns of virus-host co-divergence, particularly in lower vertebrates, which is consistent with previous findings (Geoghegan et al. 2017; Hartlage et al. 2016; Porter et al. 2020; Shi et al. 2018). The results of our cophylogenetic analysis are certainly influenced by the sample of virus diversity and will likely change as more are identified.

Wenling moray eel hepacivirus (AVM87555) forms a sister group to the ‘pegi-hepaci’ lineage-although this may be artifactual, due to recombination or extreme rate variation (Porter et al. 2020). If the position of Wenling moray eel hepacivirus is correct, this suggests that a common ancestor of the ‘pegi-hepaci’ lineage may have existed in an aquatic environment. This notion is supported by the recent finding of ‘pegi-hepaci’ derived EVE in a marine mollusc (Bamford et al. 2022). The apparent lack of pegiviruses in aquatic vertebrate and invertebrate species in this study does not equate to their absence in these organisms due to the current depth of SRA libraries available.

A key observation from this study was the identification of a flavivirus in non-bilaterians, which raises additional questions on the ancestral mode of flavivirus transmission. Non-bilaterians lack the circulatory system of vertebrates, suggesting that an alternative mode of cell-to-cell virus transmission may exists in these animals, for example cell-to-cell via exosomes (Bamford et al. (2022) as previously reported in tick-borne flaviviruses (Zhou et al. 2018).

In sum, through a broad-scale survey of publicly available transcriptomes, we uncover a wide diversity of flaviviral sequences in undersampled metazoan species. In doing so, we provide evidence for the ancient origin of the flaviviruses, closer to the emergence of the metazoans some 750–800 million years ago, and for the long-term association between the *Flaviviridae* and the Metazoa as a whole.

## Data availability

All tunicate sequence reads are available on the NCBI Sequence Read Archive (SRA) under BioProject XXXX. All viral genomes and corresponding sequences assembled in this study will be deposited in the NCBI GenBank upon acceptance. The sequences, alignments, and phylogenetic trees generated in this study will be available at https://github.com/JonathonMifsud/Transcriptome_mining_extends_the_host_range_of_the_Flaviviridae_to_non-bilaterians

## Acknowledgements

We acknowledge the University of Sydney’s high-performance computing cluster Artemis for providing the computing resources used for this study. This work would not be possible without the sequencing data that has been generously shared and we are grateful to the NCBI SRA team for their management of this important resource. We also thank A.H. McGrath for assistance in field sampling.

