## Supplemental Figures for "Transcriptome mining extends the host range of the *Flaviviridae* to non-bilaterians"

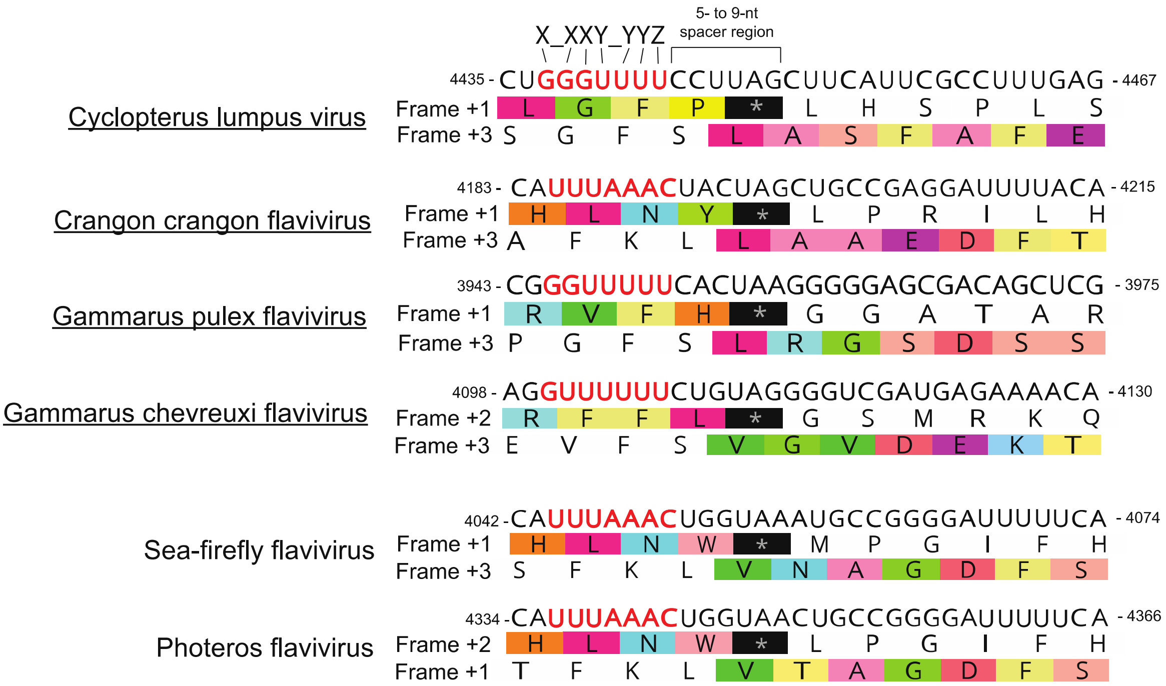


**Supplementary Figure 1.** Alignment of the predicted structured programmed ribosomal frameshift (−1 PRF) regions identified with reference sequences. Slippery heptanucleotide regions are highlighted in red. Predicted translated protein regions are indicated in colour.

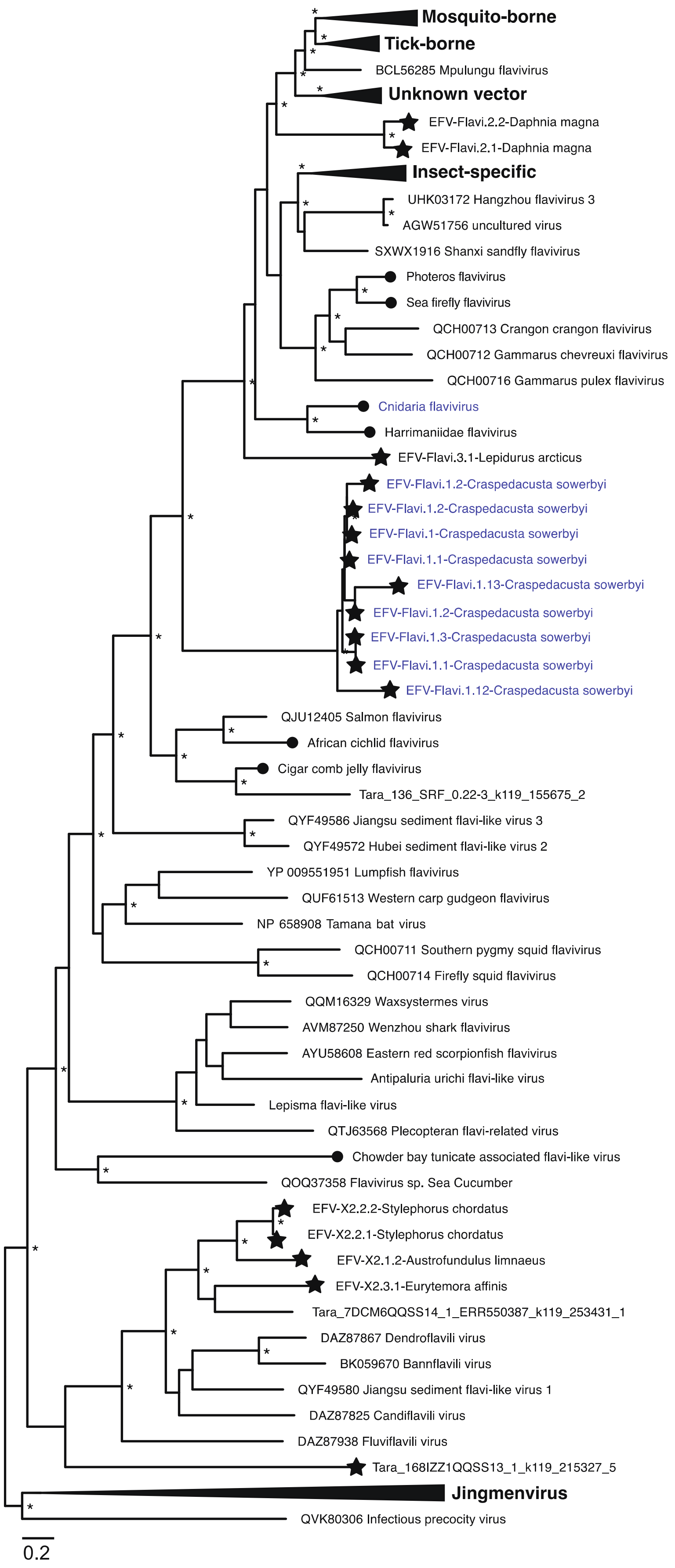

Mosquito-borne
Tick-borne
BCL56285 Mpulungu flavivirus
Unknown vector
EFV-Flavi.2.2-Daphnia magna
EFV-Flavi.2.1-Daphnia magna
Insect-specific
UHK03172 Hangzhou flavivirus 3
AGW51756 uncultured virus
SXWX1916 Shanxi sandfly flavivirus
Photeros flavivirus
Sea firefly flavivirus
QCH00713 Crangon crangon flavivirus
QCH00712 Gammarus chevreuxi flavivirus
QCH00716 Gammarus pulex flavivirus
Cnidaria flavivirus
Harrimaniidae flavivirus
EFV-Flavi.3.1-Lepidurus arcticus
EFV-Flavi.1.2-Craspedacusta sowerbyi
EFV-Flavi.1.2-Craspedacusta sowerbyi
EFV-Flavi.1-Craspedacusta sowerbyi
EFV-Flavi.1.1-Craspedacusta sowerbyi
EFV-Flavi.1.13-Craspedacusta sowerbyi
EFV-Flavi.1.2-Craspedacusta sowerbyi
EFV-Flavi.1.3-Craspedacusta sowerbyi
EFV-Flavi.1.1-Craspedacusta sowerbyi
EFV-Flavi.1.12-Craspedacusta sowerbyi
QJU12405 Salmon flavivirus
African cichlid flavivirus
Cigar comb jelly flavivirus
Tara_136_SRF_0.22-3_k119_155675_2
QYF49586 Jiangsu sediment flavi-like virus 3
QYF49572 Hubei sediment flavi-like virus 2
YP 009551951 Lumpfish flavivirus
QUF61513 Western carp gudgeon flavivirus
NP 658908 Tamana bat virus
QCH00711 Southern pygmy squid flavivirus
QCH00714 Firefly squid flavivirus
QQM16329 Waxsystermes virus
AVM87250 Wenzhou shark flavivirus
AYU58608 Eastern red scorpionfish flavivirus
Antipaluria urichi flavi-like virus
Lepisma flavi-like virus
QTJ63568 Plecopteran flavi-related virus
Chowder bay tunicate associated flavi-like virus
QOQ37358 Flavivirus sp. Sea Cucumber
EFV-X2.2.2-Stylephorus chordatus
EFV-X2.2.1-Stylephorus chordatus
EFV-X2.1.2-Austrofundulus limnaeus
EFV-X2.3.1-Eurytemora affinis
Tara_7DCM6QQSS14_1_ERR550387_k119_253431_1
DAZ87867 Dendroflavili virus
BK059670 Bannflavili virus
QYF49580 Jiangsu sediment flavi-like virus 1
DAZ87825 Candiflavili virus
DAZ87938 Fluviflavili virus
Tara_168IZZ1QQSS13_1_k119_215327_5
Jingmenvirus
QVK80306 Infectious precocity virus
0.2

**Supplementary Figure 2. Phylogenetic relationships of the flavi-like viruses and flavivirus-derived EVEs.** Phylogenetic relationships of the ‘flavi-jingmen’ clade including the EVE identified in Bamford et al. (2022) (black stars). ML phylogenetic trees based on the conserved amino acid in the NS5 region show the topological position of virus-like sequences discovered in this study (black circles) in the context of their closest relatives. All branches are scaled to the number of amino acid substitutions per site, and trees were midpoint rooted for clarity only. An asterisk indicates node support where SH-aLRT >= 80% and UFboot >= 95%.

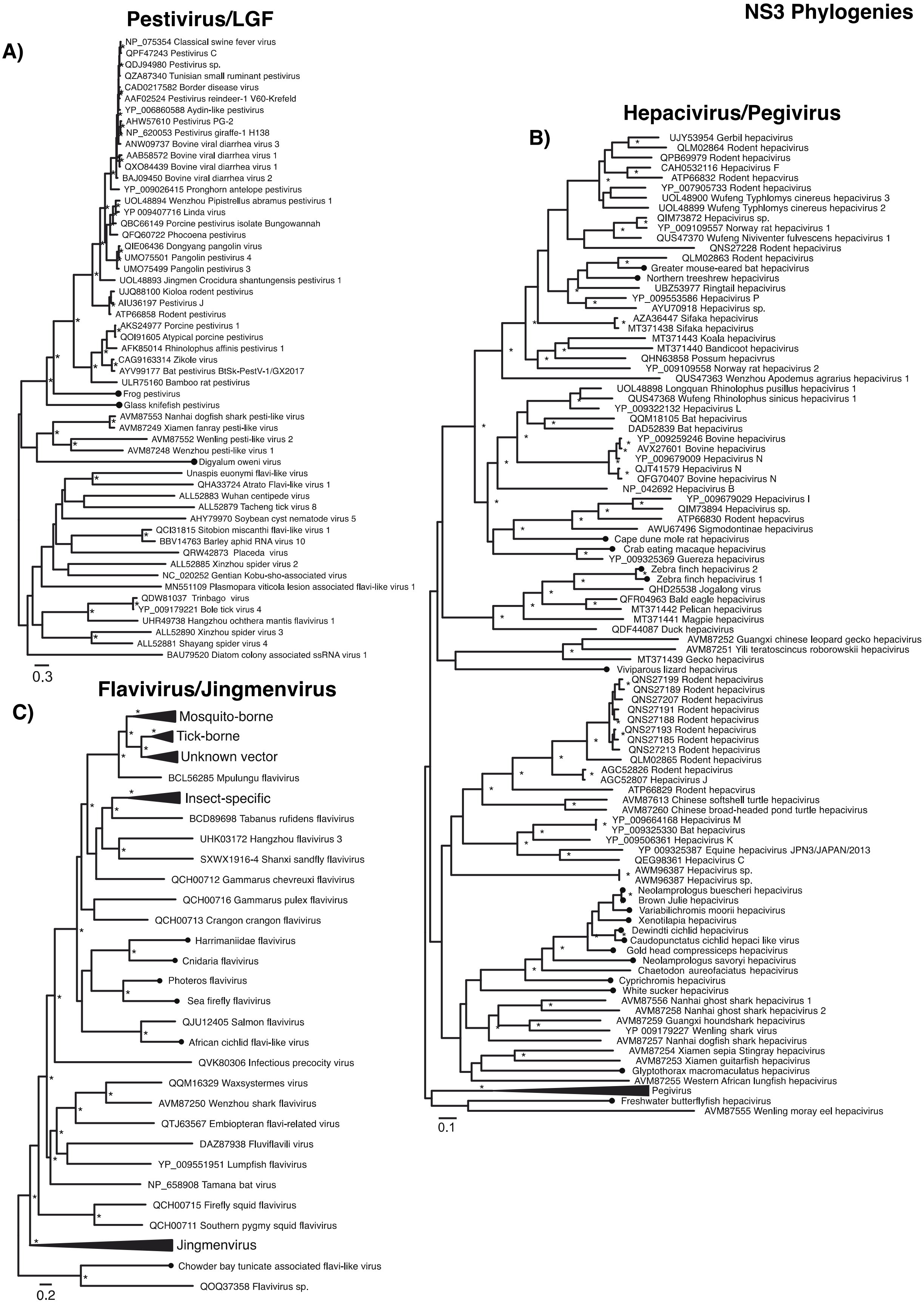
NS3 Phylogenies
Pestivirus/LGF
A)
NP_075354 Classical swine fever virus
QPF47243 Pestivirus C
QDJ94980 Pestivirus sp.
QZA87340 Tunisian small ruminant pestivirus
CAD0217582 Border disease virus
AAF02524 Pestivirus reindeer-1 V60-Krefeld
YP_006860588 Aydin-like pestivirus
AHW57610 Pestivirus PG-2
NP_620053 Pestivirus giraffe-1 H138
ANW09737 Bovine viral diarrhea virus 3
AAB58572 Bovine viral diarrhea virus 1
QXO84439 Bovine viral diarrhea virus 1
BAJ09450 Bovine viral diarrhea virus 2
YP_009026415 Pronghorn antelope pestivirus
UOL48894 Wenzhou Pipistrellus abramus pestivirus 1
YP 009407716 Linda virus
QBC66149 Porcine pestivirus isolate Bungowannah
QFQ60722 Phocoena pestivirus
QIE06436 Dongyang pangolin virus
UMO75501 Pangolin pestivirus 4
UMO75499 Pangolin pestivirus 3
UOL48893 Jingmen Crocidura shantungensis pestivirus 1
UJQ88100 Kioloa rodent pestivirus
AIU36197 Pestivirus J
ATP66858 Rodent pestivirus
AKS24977 Porcine pestivirus 1
QOI91605 Atypical porcine pestivirus
AFK85014 Rhinolophus affinis pestivirus 1
CAG9163314 Zikole virus
AYV99177 Bat pestivirus BtSk-PestV-1/GX2017
ULR75160 Bamboo rat pestivirus
Frog pestivirus
Glass knifefish pestivirus
AVM87553 Nanhai dogfish shark pesti-like virus
AVM87249 Xiamen fanray pesti-like virus
AVM87552 Wenling pesti-like virus 2
AVM87248 Wenzhou pesti-like virus 1
Digyalum oweni virus
Unaspis euonymi flavi-like virus
QHA33724 Atrato Flavi-like virus 1
ALL52883 Wuhan centipede virus
ALL52879 Tacheng tick virus 8
AHY79970 Soybean cyst nematode virus 5
QCI31815 Sitobion miscanthi flavi-like virus 1
BBV14763 Barley aphid RNA virus 10
QRW42873 Placeda virus
ALL52885 Xinzhou spider virus 2
NC_020252 Gentian Kobu-sho-associated virus
MN551109 Plasmopara viticola lesion associated flavi-like virus 1
QDW81037 Trinbago virus
YP_009179221 Bole tick virus 4
UHR49738 Hangzhou ochthera mantis flavivirus 1
ALL52890 Xinzhou spider virus 3
ALL52881 Shayang spider virus 4
BAU79520 Diatom colony associated ssRNA virus 1
0.3
Flavivirus/Jingmenvirus
C)
Mosquito-borne
Tick-borne
Unknown vector
BCL56285 Mpulungu flavivirus
Insect-specific
BCD89698 Tabanus rufidens flavivirus
UHK03172 Hangzhou flavivirus 3
SXWX1916-4 Shanxi sandfly flavivirus
QCH00712 Gammarus chevreuxi flavivirus
QCH00716 Gammarus pulex flavivirus
QCH00713 Crangon crangon flavivirus
Harrimaniidae flavivirus
Cnidaria flavivirus
Photeros flavivirus
Sea firefly flavivirus
QJU12405 Salmon flavivirus
African cichlid flavi-like virus
QVK80306 Infectious precocity virus
QQM16329 Waxsystermes virus
AVM87250 Wenzhou shark flavivirus
QTJ63567 Embiopteran flavi-related virus
DAZ87938 Fluviflavili virus
YP_009551951 Lumpfish flavivirus
NP_658908 Tamana bat virus
QCH00715 Firefly squid flavivirus
QCH00711 Southern pygmy squid flavivirus
Jingmenvirus
Chowder bay tunicate associated flavi-like virus
QOQ37358 Flavivirus sp.
0.2
Hepacivirus/Pegivirus
B)
UJY53954 Gerbil hepacivirus
QLM02864 Rodent hepacivirus
QPB69979 Rodent hepacivirus
CAH0532116 Hepacivirus F
ATP66832 Rodent hepacivirus
YP_007905733 Rodent hepacivirus
UOL48900 Wufeng Typhlomys cinereus hepacivirus 3
UOL48899 Wufeng Typhlomys cinereus hepacivirus 2
QIM73872 Hepacivirus sp.
YP_009109557 Norway rat hepacivirus 1
QUS47370 Wufeng Niviventer fulvescens hepacivirus 1
QNS27228 Rodent hepacivirus
QLM02863 Rodent hepacivirus
Greater mouse-eared bat hepacivirus
Northern treeshrew hepacivirus
UBZ53977 Ringtail hepacivirus
YP_009553586 Hepacivirus P
AYU70918 Hepacivirus sp.
AZA36447 Sifaka hepacivirus
MT371438 Sifaka hepacivirus
MT371443 Koala hepacivirus
MT371440 Bandicoot hepacivirus
QHN63858 Possum hepacivirus
YP_009109558 Norway rat hepacivirus 2
QUS47363 Wenzhou Apodemus agrarius hepacivirus 1
UOL48898 Longquan Rhinolophus pusillus hepacivirus 1
QUS47368 Wufeng Rhinolophus sinicus hepacivirus 1
YP_009322132 Hepacivirus L
QQM18105 Bat hepacivirus
DAD52839 Bat hepacivirus
YP_009259246 Bovine hepacivirus
AVX27601 Bovine hepacivirus
YP_009679009 Hepacivirus N
QJT41579 Hepacivirus N
QFG70407 Bovine hepacivirus N
NP_042692 Hepacivirus B
YP_009679029 Hepacivirus I
QIM73894 Hepacivirus sp.
ATP66830 Rodent hepacvirus
AWU67496 Sigmodontinae hepacivirus
Cape dune mole rat hepacivirus
Crab eating macaque hepacivirus
YP_009325369 Guereza hepacivirus
Zebra finch hepacivirus 2
Zebra finch hepacivirus 1
QHD25538 Jogalong virus
QFR04963 Bald eagle hepacivirus
MT371442 Pelican hepacivirus
MT371441 Magpie hepacivirus
QDF44087 Duck hepacivirus
AVM87252 Guangxi chinese leopard gecko hepacivirus
AVM87251 Yili teratoscincus roborowskii hepacivirus
MT371439 Gecko hepacivirus
Viviparous lizard hepacivirus
QNS27199 Rodent hepacivirus
QNS27189 Rodent hepacivirus
QNS27207 Rodent hepacivirus
QNS27191 Rodent hepacivirus
QNS27188 Rodent hepacivirus
QNS27193 Rodent hepacivirus
QNS27185 Rodent hepacivirus
QNS27213 Rodent hepacivirus
QLM02865 Rodent hepacivirus
AGC52826 Rodent hepacivirus
AGC52807 Hepacivirus J
ATP66829 Rodent hepacvirus
AVM87613 Chinese softshell turtle hepacivirus
AVM87260 Chinese broad-headed pond turtle hepacivirus
YP_009664168 Hepacivirus M
YP_009325330 Bat hepacivirus
YP_009506361 Hepacivirus K
YP 009325387 Equine hepacivirus JPN3/JAPAN/2013
QEG98361 Hepacivirus C
AWM96387 Hepacivirus sp.
AWM96387 Hepacivirus sp.
Neolamprologus buescheri hepacivirus
Brown Julie hepacivirus
Variabilichromis moorii hepacivirus
Xenotilapia hepacivirus
Dewindti cichlid hepacivirus
Caudopunctatus cichlid hepaci like virus
Gold head compressiceps hepacivirus
Neolamprologus savoryi hepacivirus
Chaetodon aureofaciatus hepacivirus
Cyprichromis hepacivirus
White sucker hepacivirus
AVM87556 Nanhai ghost shark hepacivirus 1
AVM87258 Nanhai ghost shark hepacivirus 2
AVM87259 Guangxi houndshark hepacivirus
YP 009179227 Wenling shark virus
AVM87257 Nanhai dogfish shark hepacivirus
AVM87254 Xiamen sepia Stingray hepacivirus
AVM87253 Xiamen guitarfish hepacivirus
Glyptothorax macromaculatus hepacivirus
AVM87255 Western African lungfish hepacivirus
Pegivirus
Freshwater butterflyfish hepacivirus
AVM87555 Wenling moray eel hepacivirus
0.1

**Supplementary Figure 3. Alternative phylograms based on the conserved NS3 region.** Phylogenetic relationships of the ‘flavi-jingmen’, ‘pegi-hepaci’, and ‘pesti-LGF’ clades. ML phylogenetic trees based on the conserved amino acid in the NS3 region show the topological position of virus-like sequences discovered in this study (black circles) in the context of their closest relatives. All branches are scaled to the number of amino acid substitutions per site, and trees were midpoint rooted for clarity only. An asterisk indicates node support where SH-aLRT >= 80% and UFboot >= 95%.

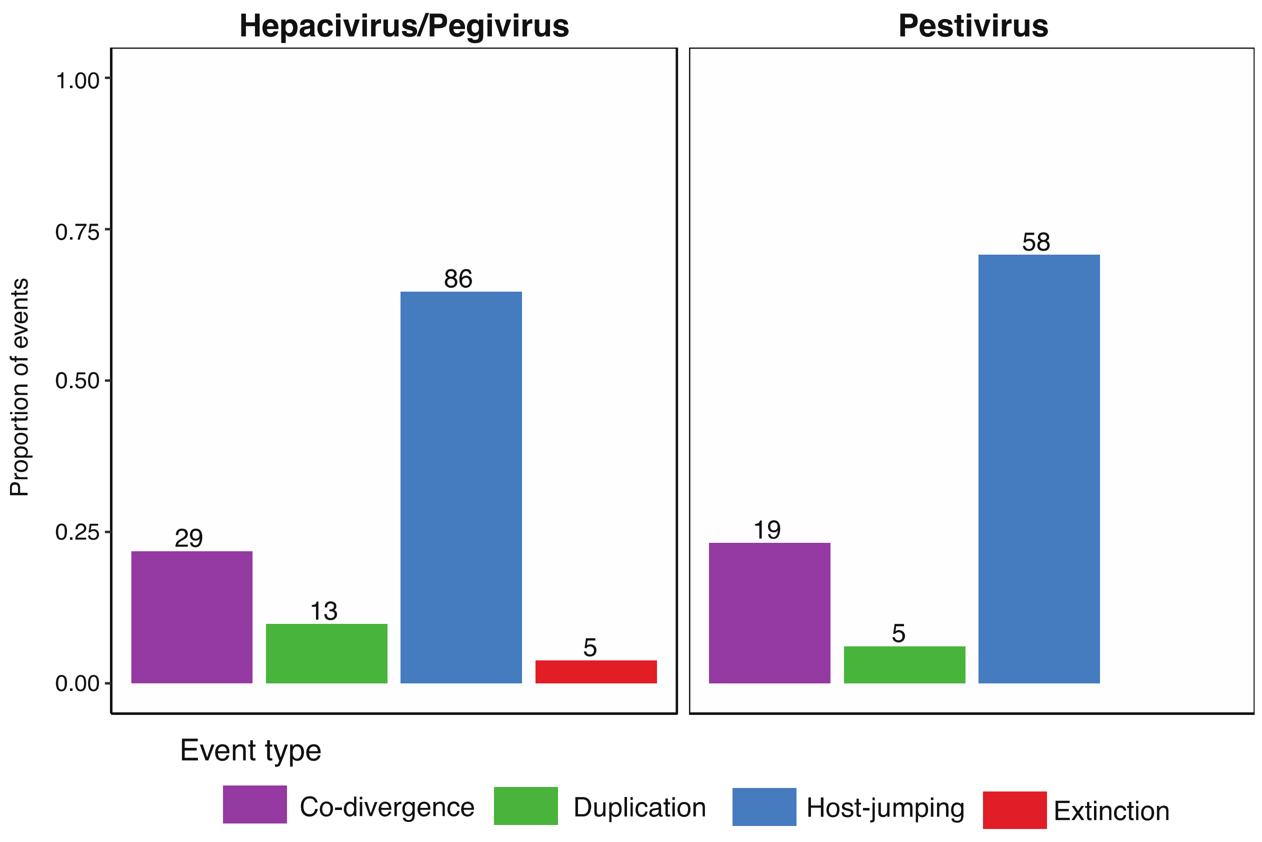


**Supplementary Figure 4. Reconciliation analysis**. Reconciliation analysis of select virus groups. Bar plots illustrate the range of the proportion of possible events and are coloured by event type.

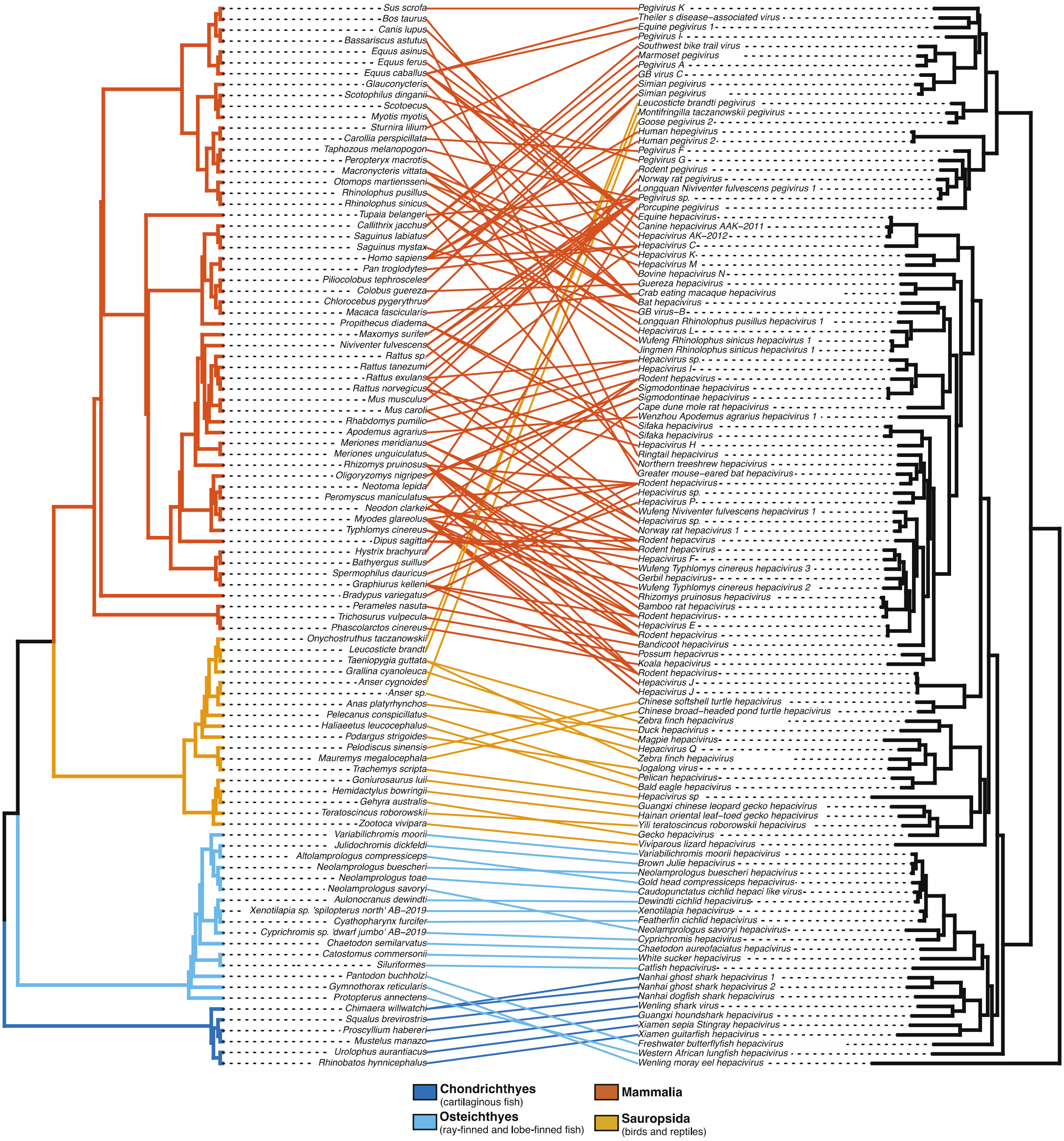


**Supplementary Figure 5. Tanglegram of the 'pegi-hepaci' clade and their hosts with species labels present**. Branches of the host tree (left) and lines are colored to represent the host clade. All branches on the virus tree are scaled to the number of amino acid substitutions per site, and both trees were midpoint rooted for clarity only.
